# External Microbiota of Western United States Bats: Does It Matter Where You Are From?

**DOI:** 10.1101/017319

**Authors:** A.K. Kooser, J.C. Kimble, J.M. Young, D.C. Buecher, E.W. Valdez, A. Porras-Alfaro, D.E. Northup

## Abstract

White-nose syndrome (WNS), a disease caused by the fungus *Pseudogymnoascus destructans*^1^, has spread west from New York to Missouri and has killed more than six million bats^2^. In bat hibernacula where WNS is present, mass mortality has been observed and there is a high potential for population collapse or extinction of some species at a regional level. Although WNS is not yet present in the western U.S., the high diversity ofbat species^3^ and appropriate conditions for *P. destructans* in area caves may put these populations at risk. The absence of WNS in western caves provides a unique opportunity to ask questions about how bat species, geographic location, and habitat shape pre-WNS bat microbiota. The importance of microbiota is shown in many organisms, including amphibians, where individuals that survive a chytrid infection carry a higher prevalence of *Janthinobacterium lividum*^4^. The establishment of a pre-WNS baseline microbiota of western bats is critical to understanding how *P. destructans* may impact the native microbiota of the bats. Previous studies^5^^,^^6^ that identified the microbiota of bats have focused on gut and fecal microbiota, with little attention given to the external microbiota. Here we show for the first time that habitat and geography influence differences in the abundance and diversity of external bat microbiota. From our 202 (62 cave-netted, 140 surface-netted) bat samples belonging to 13 species of western bats uninfected with WNS, we identified differences in microbiota diversity among sites, and between cave-netted versus surface-netted bats, regardless of sex and species. These results present novel information about the factors that shape external microbiota of bats providing new insights into patterns of diversity in a pre-WNS bat population.

---

Since the discovery of WNS in 2006-2007 in New York^7^, there is conclusive evidence indicating that *Pseudogymnoascus destructans* acts as the primary pathogen causing the disease. Moreover, through controlled experiments, it was determined that WNS is spread by direct contact with this fungus^8^. *P. destructans* is often fatal to bats because of its ability to colonize and penetrate bat tissue, which disrupts hibernation by causing frequent arousals of bats in hibernacula^9^^,^^10^. Consequently, this leads to depletion of critical energy reserves stored as fat and an inability to maintain water homeostasis^11^. Research evidence suggests that *P. destructans* originated in Europe, accounting for the absence of mass mortality of bats in European hibernacula infected with *P. destructans*^12^.

The westward movement of WNS is on a trajectory that will allow it to enter the West through Colorado and New Mexico’s respective southern and northern borders. Within these regions are western analogs (e.g., *Myotis evotis*) of bat species that have been greatly impacted by WNS in the east (e.g., *Myotis septentrionalis*) and would likely succumb to the same fate. Potentially, over 16 western bat species could be affected by this disease. Thus, given the rapid westward spread of WNS and our limited knowledge about the susceptibility of western bat populations, there is a need to establish the baseline microbiota across key western bat species. This pre-WNS external microbiota dataset of western U.S. bat populations will serve as a resource for future studies that investigate the differences in vulnerabilities of different bat species, as well as aid in identifying the dynamics that influence the occurrence of microbial communities present on the surface of bats. Similar to gut microbiota^13^, external microbiota^14^ may suppress external bacterial and fungal infections, as seen with the chytrid fungal infections in amphibians^15^. The microbiota patterns documented in our study will provide insight into the diversity of a pre-WNS bat population across states that have habitats that are vulnerable to WNS. We hypothesize that habitat, geography, and bat clustering behavior account for the diversity of external microbiota communities found on bats.

To address our hypothesis, during the springs and summers of 2013 and 2014, we collected a pre-WNS dataset of 202 western bats from five locations in the Southwest, including Carlsbad Caverns National Park (CCNP), Fort Stanton-Snowy River Cave National Conservation Area (FS), El Malpais National Monument (ELMA), and caves near Roswell (HGL), New Mexico, as well as Grand Canyon Parashant National Monument (PARA), Arizona, to determine which factors influence the bat microbiota. We characterized samples by geographic sites (captured bats between and within New Mexico and Arizona, Figure 1), cave versus surface-netted bats, sex, and species (Supplemental Data 1). Analysis of the data by sex and species found weak to no evidence (ratio of random error was 2 or less) as predictors of microbiota composition; therefore, our metadata analysis focused on sites of capture and cave versus surface-netted samples. Alpha diversity indices of bacterial and fungal samples (Figure 2a, b, respectively) show differences in mean diversity between sites and a large variation of diversity within a site. The highest observed bacterial diversity was seen in bats sampled at ELMA. The lowest observed bacterial diversity was from HGL. For fungal samples, the highest observed diversity was from FS and lowest from HGL. Across all samples the observed fungal diversity was much lower than the bacterial diversity.

**Figure 1.**
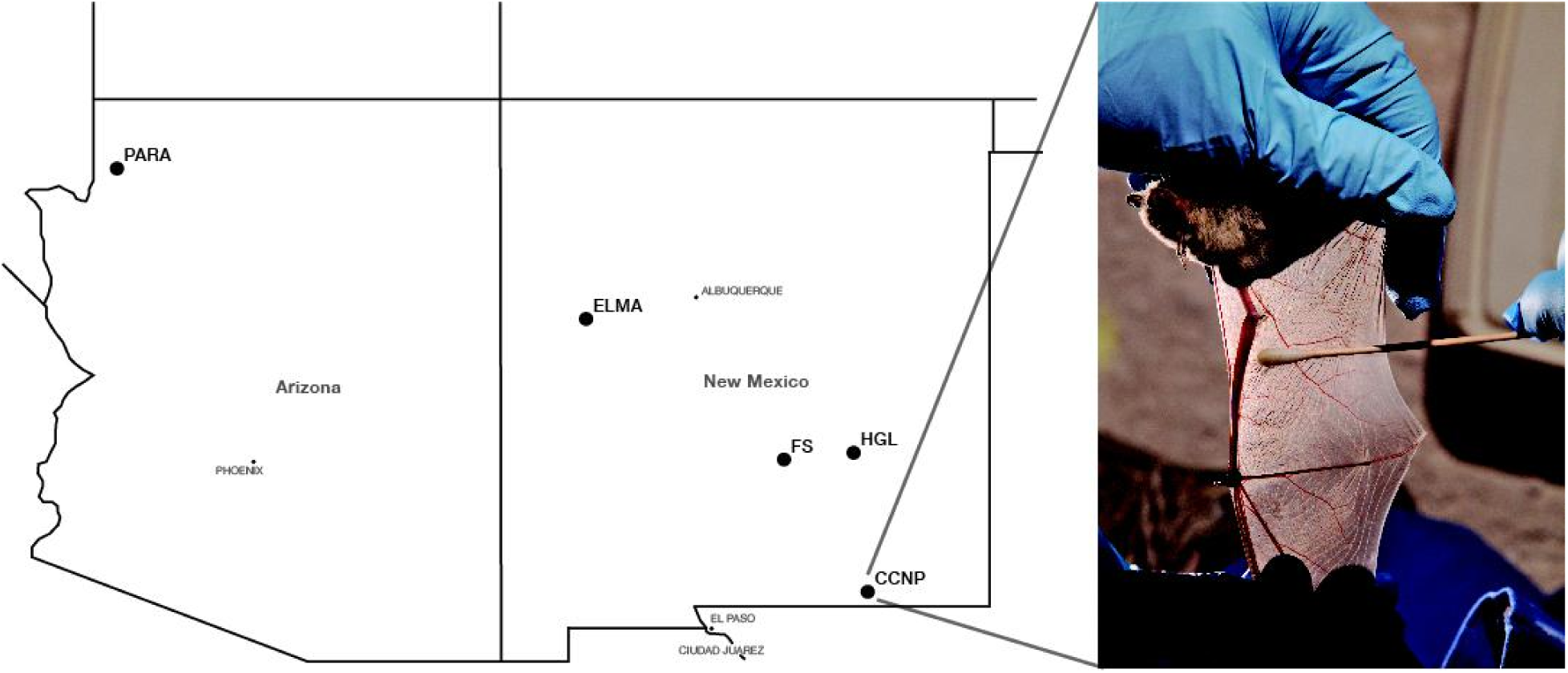
Location of field sites and example swabbing. **a,** Map of the field sites where bats were collected for this study (Map designed by Ara Kooser CC-BY 3.0. Data by OpenStreetMap, under ODbL) and **b,** Swabbing of a cave bat (*Myotis velifer*) netted in Left Hand Tunnel, CAVE for genomic DNA.

**Figure 2.**
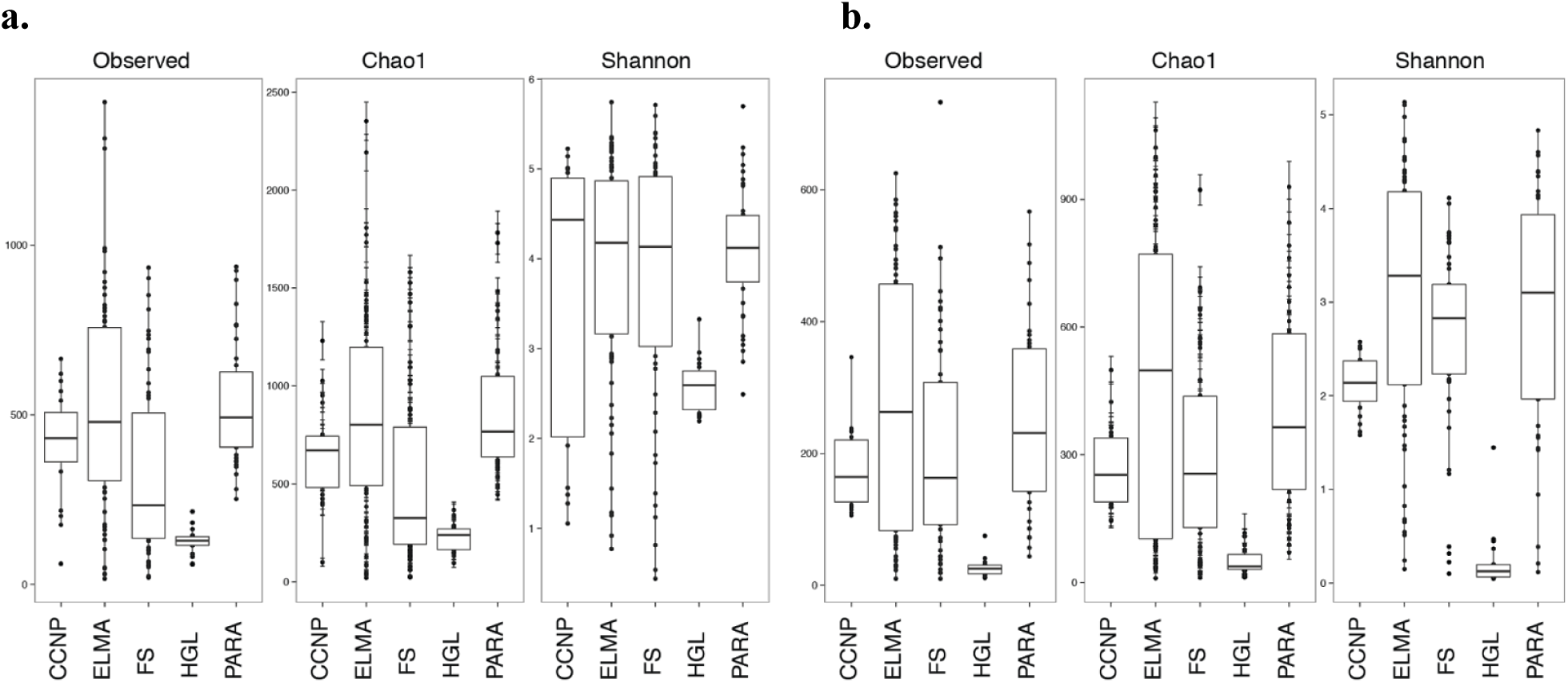
Observed diversity of microbiota across regions for El Malpais National Monument (ELMA), Fort Stanton-Snowy River Cave National Conservation Area (FS), Grand Canyon Parashant National Monument (PARA), Carlsbad Caverns National Park (CCNP), and High Grasslands (HGL). **a,** Box plot of alpha diversity indices for microbial communities separated by region and **b,** Box plot of alpha diversity indices for fungal communities separated by region.

Non-metric multidimensional scaling plots of bacterial and fungal communities by site (i.e., ELMA, HGL, FS, CCNP, and PARA) show five groupings (Figures 3a, b). The HGL and PARA samples form two distinct tight clusters, while the remaining groups form looser associations. The NMDS plots of bacteria and fungi from cave and surface-netted bats show two distinctive groups with little overlap for the bacterial samples (Figures 4a, b). To further test if the observed patterns in the NMDS could be attributed to habitat and geography parameters, we used a random forest model. We tested whether microbiota composition could identify samples based on cave versus surface-netted and by site. The ratio of random error was 3.63 among bacteria samples from all sites, whereas the ratio of random error was 6.25 between cave versus surface-netted bacteria samples. The ratio of random error for fungal samples among all sites was 5.23 and for cave versus surface-netted it was 2.56. Cave versus surface-netted was the most predictive for bacterial samples, while site was most predictive for fungal samples.

**Figure 3.**
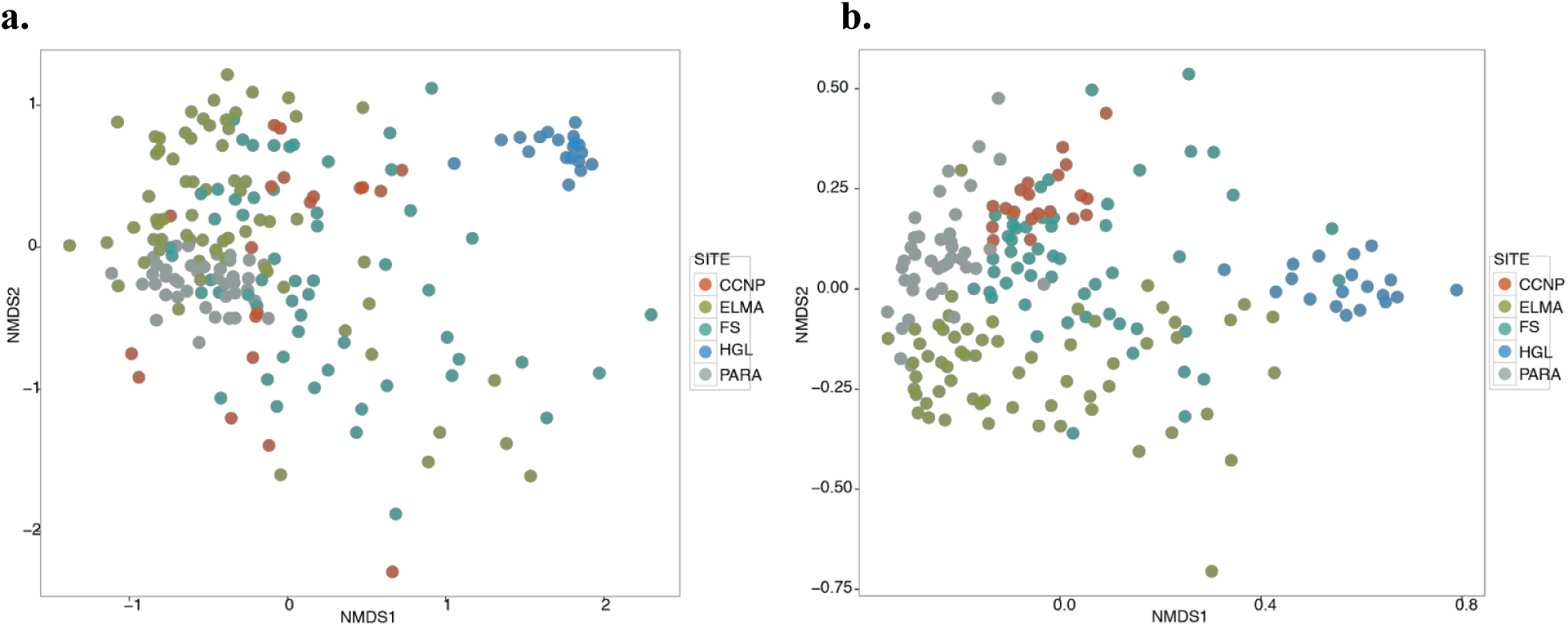
Regional relationships between microbiota communities and biogeographic parameters for El Malpais National Monument (ELMA), Fort Stanton-Snowy River Cave National Conservation Area (FS), Grand Canyon Parashant National Monument (PARA), Carlsbad Caverns National Park (CCNP), and High Grasslands (HGL). **a,** Bacterial NMDS plot colored by area the bats were caught **b,** Fungal NMDS plot colored by area the bats were caught.

**Figure 4.**
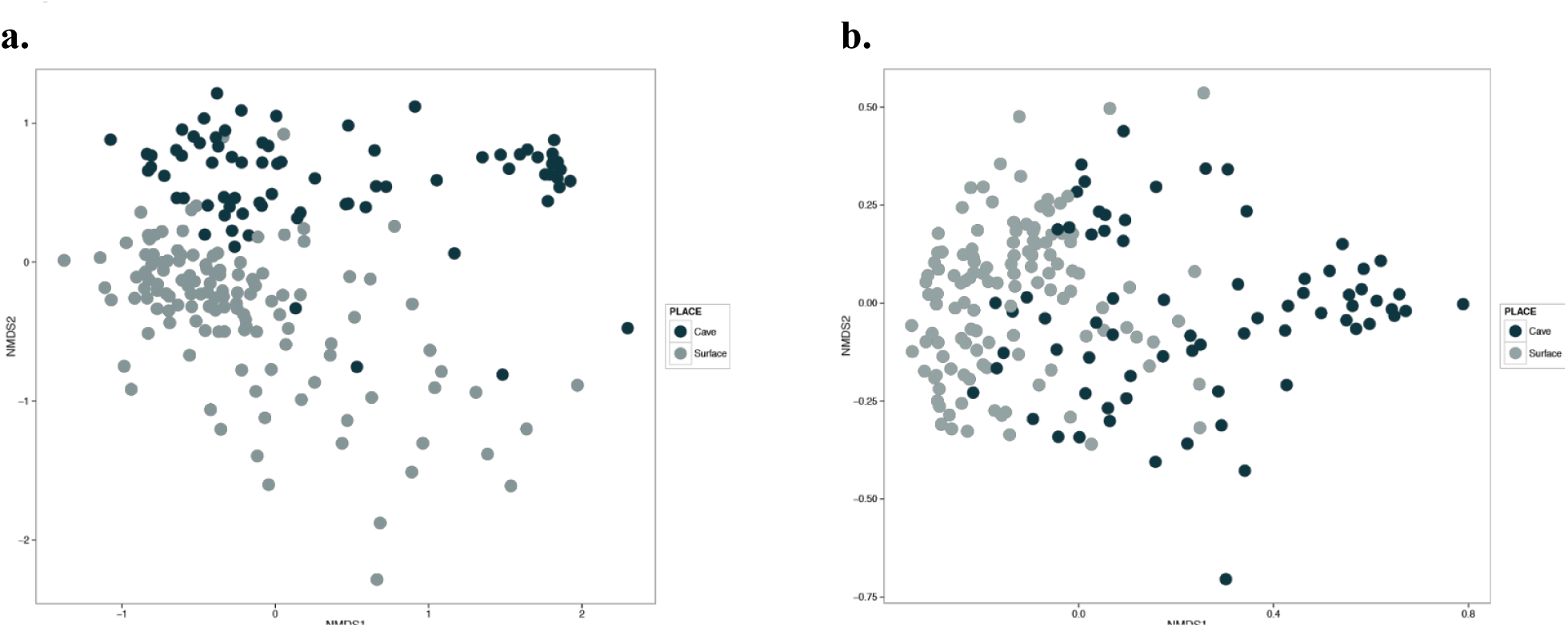
Cave versus surface-netted relationships between microbiota communities and cave versus surfaced netted bats microbiota for El Malpais National Monument (ELMA), Fort Stanton-Snowy River Cave National Conservation Area (FS), Grand Canyon Parashant National Monument (PARA), Carlsbad Caverns National Park (CCNP), and High Grasslands (HGL) **a,** Bacterial NMDS plot colored by cave or surface netted **b,** Fungal NMDS plot colored by cave or surface-netted.

The significance of habitat and geography on community structure was tested using a poisson-lognormal generalized linear mixed model analysis of count data. Operational taxonomic units (OTUs) that show up as significant at a false discovery rate correction <0.05 for bacterial phyla (Extended Data Figure 1) include Acidobacteria, Actinobacteria, Alphaproteobacteria, Bacteroidetes, Betaproteobacteria, Chloroflexi, Cyanobacteria, Firmicutes, Gemmatimonadetes, Tenericutes, and TM7 among all sites. Between cave versus surface-netted bats, the significant bacterial phyla include Acidobacteria, Actinobacteria, Alphaproteobacteria, Armatimonadetes, Chloroflexi, Cyanobacteria, Deltaproteobacteria, Firmicutes, Nitrospirae, Synergistetes, and Thermi. In the fungal communities, the following classes (Extended Data Figure 2) differed by site: Ascomycota unidentified, Dothideomycetes, Eurotiomycetes, Lecanoromycetes, Leotiomycetesm Pezizomycetes, Saccharomycetes, Sordariomycetes, Agaricomycetes, Tremellomycetes, and Incertae sedis. Dothideomycetes, Eurotiomycetes, Leotiomycetes, Pezizomycetes, Saccharomycetes, Ascomycota unidentified, and Agaricomycetes were the classes that differed between cave and surface-netted bats.

Our results show a complex relationship of external bat microbiota with respect to habitat and geographic effects such as site of capture and habitat (cave versus surface-netted). We concluded that species or sex of bat was not significant in determining microbiota composition. However, geographic site, and habitat are significant in determining diversity observed for both bacterial and fungal microbiota. One explanation for the minimal overlap on the NMDS of the microbiota found on bats from cave versus surface-netted sites may be the result of microbiota community turnover as the bats opportunistically change roosting areas. This turnover is hypothesized to be similar to what has been observed in the variation in human hand microbiota from interactions with household surfaces^15^. We attribute differences in microbial alpha diversity to microbiota acquired in the habitats in which bats were captured. Additionally, we believe low diversity of bacteria and fungal microbiota in HGL is related to having a less complex landscape in the form of a sparse grassland habitat, fewer social interactions and clustering behavior of the single bat species present in these caves.

Our discovery of habitat and geographic patterns related to the external microbiota of western bats shows that bat behavior and local roosting habitat drive the patterns in microbiota diversity. We suggest future investigations should include a broad range of habitat types and associated bat species along differing latitudinal and longitudinal gradients to better understand the observed patterns in diversity. Fundamental questions should be addressed, such as, “Does specific site location within a geographic area (e.g., Colorado Plateau) or sampling locality (e.g., cave and surface), as well as the number of species occupying these sites during sampling and time of year, affect the composition of external microbiota?” Additionally, future findings can provide insight into the microbial community relationships between regions where *P. destructans* is present, with and without WNS, and which natural occurring bat bacteria and fungi can be used to suppress WNS.

## Methods

We sampled 202 bats (62 cave and 104 surface-netted), belonging to 13 species (Supplemental Data 1), for external microbiota identification from a total of five study sites in the Southwest: Grand Canyon Parashant National Monument (PARA), in Arizona, and Carlsbad Caverns National Park (CCNP), Fort Stanton (FS), El Malpais National Monument (ELMA), and Bureau of Land Management (HGL) caves near Roswell, New Mexico. Bat sample collection was allowed under the following permits: 2014 Arizona and New Mexico Game and Fish Department Scientific Collecting Permit (SP670210, SCI#3423, SCI#3350), National Park Service Scientific Collecting Permit (CAVE-2014-SCI-0012, ELMA-2013-SCI-0005, ELMA-2014-SCI-0001, PARA-2012-SCI-0003), Fort Collins Science Center Standard Operating Procedure (SOP) SOP#: 2013-01, and an Institutional Animal Care and Use Committee (IACUC) Permit from the University of New Mexico (Protocol #12-100835-MCC) and from the National Park Service (Protocol #IMR_ELMA.PARA_Northup_Bat_2013.A2). The skin (i.e., ears, wings and uropatagia) and furred surfaces of the bat were swabbed (Figure 1b) with sterile swabs soaked in sterile Ringer’s solution^16^. Each swab was placed in a sterile 1.7ml snap-cap microfuge tube and immediately frozen in a liquid nitrogen dry shipper. Samples were transported to the University of New Mexico and stored in a -80°C freezer. We used MR DNA Molecular Research LP, Shallowater, Texas (http://www.mrdnalab.com/) for genomic DNA extraction and 454 sequencing diversity assays of bacterial 16S rDNA and fungal ITS genes.

All 454 reads were processed in QIIME^17^. Bacterial sequences shorter than 200 bp or longer than 500 bp and with a quality score lower than 30 were eliminated. Bacterial samples were denoised and clustered with sumaclust^18^ and chimera checked using usearch^19^. Fungal sequences were pre-processed by discarding all sequences with a quality score lower than 30. Fungal clustering was done using the open reference picking with the sumaclust option. Taxonomy was assigned using the Greengenes 13_8 core data set^20^ with uclust and the UNITE OTUs 12_11^21^ alpha data set with SortMeRNA^22^, respectively. This yielded a total of 193 bacterial 16S and fungal ITS paired samples and 9 bacteria samples with no fungal counterpart.

Variation in community structure was visualized using the phyloseq package^23^ and ggplot2^24^ in the R software package^25^. Beta diversity was analyzed using nonmetric dimensional analysis. Random forest models were run in QIIME using 10-fold cross-validation with 1,000 trees. The MCMC.otu package^26^ in R was used to quantify proportional changes in community structure between cave and surface netted.

## Acknowledgements

We thank the staff at El Malpais and Grand Canyon Parashant National Monuments, Carlsbad Caverns National Park, Bureau of Land Management, and the Fort Stanton Cave Study Project. Funding was provided by the National Park Service (CPCESU) and Western National Park Association for work in El Malpais National Monument, Carlsbad Caverns National Park, and Grand Canyon Parashant National Monument. The Bureau of Land Management and Fort Stanton Cave Study Project funded work in Fort Stanton and BLM Caves 45 and 55. Additional funding was provided by the New Mexico Game and Fish Department Share with Wildlife Program, Cave Conservancy Foundation, Eppley Foundation, National Speleological Society Rapid Response Fund, and T&E, Inc. We thank Kait Hughes for fieldwork assistance, assistance with study design and preliminary data analysis; Graham Walmsley for writing suggestions; Brennen Reece for graphic design and typographic help; and Ken of Kenneth Ingham Photography for the bat photo. Any use of trade, firm, or product names is for descriptive purposes only and does not imply endorsement by the U.S. Government.

## Author Contributions

A.S.K contributed to the bacterial data analysis, methods, results, discussion and fieldwork. J.C.K contributed to the writing, results, and discussion. J.M.Y. contributed to the fungal data analysis, methods, results, discussion, and fieldwork. E.W.V. contributed to data collection, writing, and interpretation of data in relation to bat ecology. A. P-A contributed to writing, editing, and fungal analysis. D.E.N. contributed to study design, funding acquisition, data collection, editing, and interpretation of habitat characteristics and bacterial sequencing results. D.C.B. contributed to study design, funding acquisition, data collection and discussions regarding bat ecology.

## Author Information

Reprints and permissions information is available at *TBA*. The authors declare no competing financial interests. Correspondence and requests for materials should be addressed to A.S.K. or D.E.N. R scripts, workflow, and data for this project are available at: https://github.com/bioinfonm/microBat

This information is preliminary and is subject to revision. It is being provided to meet the need for timely best science. The information is provided on the condition that neither the U.S. Geological Survey nor the U.S. Government shall be held liable for any damages resulting from the authorized or unauthorized use of the information.

**Extended Data Figure 1.**
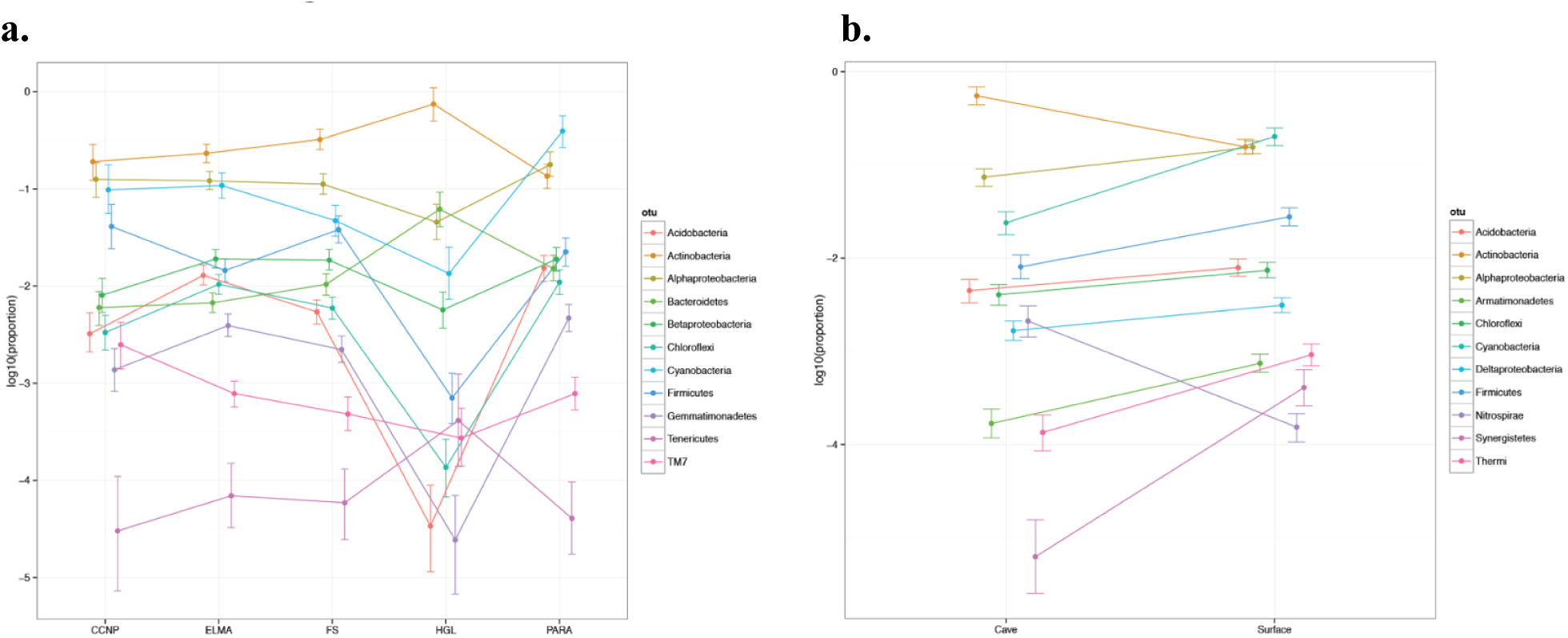

**Extended Data Figure 2.**
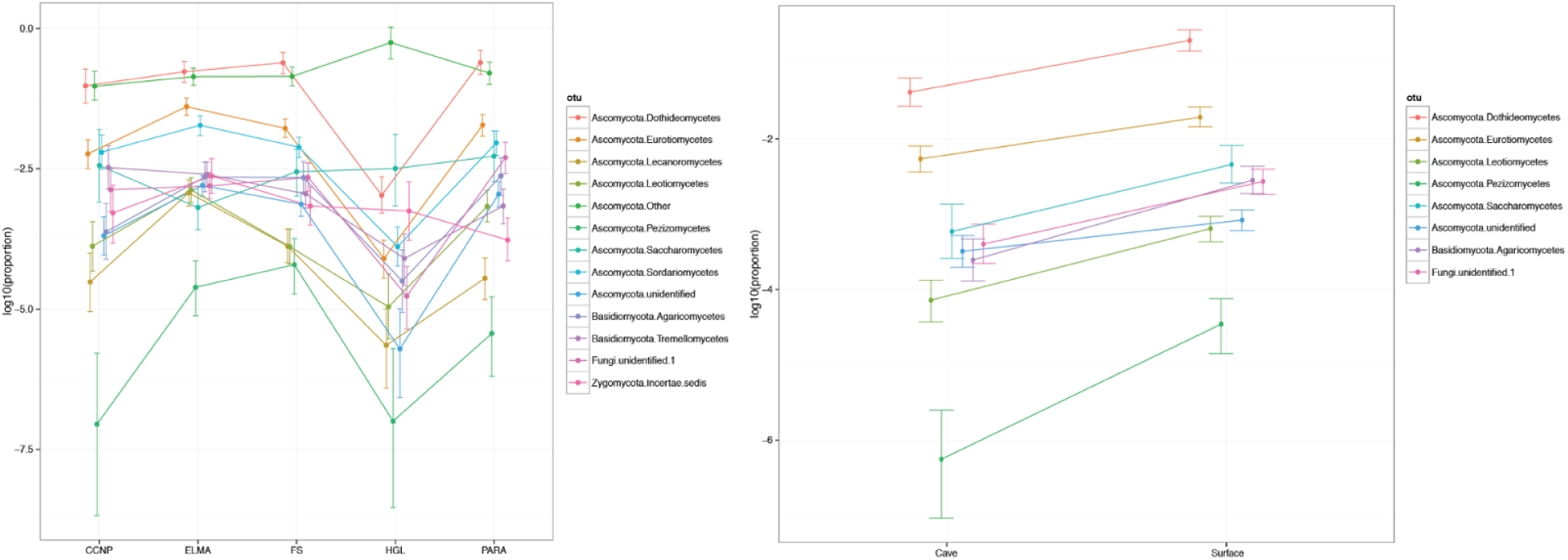

